# Interlaboratory benchmarking of differential scanning fluorimetry reveals sources of variability in protein thermal stability parameters

**DOI:** 10.64898/2026.10.12.758416

**Authors:** Sébastien Brûlé, Maelenn Chevreuil, Osvaldo Burastero, Olga Abian, Gregor Anderluh, Magali Aumont-Nicaise, Martynas Bagdonas, Holly Birchenough, Alexander K. Buell, Tatsiana Charnavets, Audrey Crousilles, Nataliya Danilenko, Markus Epe, Edoardo Fabini, Alessandra Feoli, Maria Garcia-Alai, Raphael Gasper, Erika Griss Pascualli, Josef Houser, Thomas Jowitt, Alexis Kuntzmann, Andreas Langer, Karine Lapouge, Kelvin Lau, Aline Le Roy, Blanca López-Méndez, Romain Malempré, Caroline Mas, André Matagne, Daumantas Matulis, Andre Hubertus Mueller, Sebastiano Pasqualato, Eva Paulenova, Gianluca Sbardella, Werner Streicher, Angelica Struve Garcia, Philipp O. Tsvetkov, Stephan Uebel, Marylène Vandevenne, Sonia Vega, Adrian Velazquez-Campoy, Ross William Walton, Nika Žibrat Kalanj, Patrick England, Bertrand Raynal

## Abstract

Differential scanning fluorimetry (DSF) is widely used to assess protein thermal stability and to compare protein variants, ligands, or formulations with limited sample consumption. Yet, the reproducibility of DSF-derived parameters across laboratories, instruments and analysis workflows remains poorly defined.

Here, we report a two-phase interlaboratory benchmark using a shared V_H_H sample across 26 laboratories, 28 instruments and seven DSF instrument models. In Phase 1, participants applied their routine acquisition and analysis workflows, whereas in Phase 2, acquisition conditions were harmonised. Datasets were analysed locally and then centrally reprocessed with MoltenProt.

Participant-reported apparent melting temperatures (*T*_m_) showed good within-instrument repeatability but moderate inter-instrument dispersion. Harmonisation of acquisition conditions did not substantially reduce this dispersion. Conversely, centralised reprocessing narrowed the distribution among retained datasets, indicating contributions from analysis workflows and data-acceptance criteria in addition to instrument-dependent factors. Comparison with differential scanning calorimetry (DSC) further showed that apparent *T*_m_ depended strongly on the fluorescence readout. For this V_H_H, single-wavelength fluorescence at 350 nm showed the closest agreement with the DSC transition temperature, whereas the 350/330 nm ratio and barycentric mean generally yielded higher values. Furthermore, DSF-derived apparent unfolding enthalpies were more variable than apparent *T*_m_.

Together, these findings show that DSF can be highly repeatable within a defined instrument and workflow, but that quantitative comparability across instruments depends on instrument-specific effects, the choice of fluorescence readout and analysis strategy. Improved comparability will require transparent reporting of acquisition and analysis conditions, explicit data-quality criteria, and well-characterised reference materials to monitor instrument-dependent offsets.

## Introduction

By probing protein responses to increasing temperature, thermal stability measurements provide valuable insights into conformational stability and unfolding behaviour (Privalov, 1979; Deller et al., 2016; Seelig & Seelig, 2023), with applications ranging from structural and biophysical characterisation (Boivin et al., 2013) to formulation studies (Remmele et al., 1998; Kellner et al., 2021; Lisina et al., 2023) and ligand screening (Savitski et al., 2014; Llowarch et al., 2023).

Among the available techniques to measure thermal transitions, differential scanning calorimetry (DSC) is generally viewed as the gold standard approach (Johnson, 2013; Wen et al., 2020). It directly measures the heat-capacity change of a protein solution as a function of temperature, and can provide both the unfolding temperature also referred to as the melting temperature (*T*_m_) and thermodynamic parameters associated with the transition (Freire, 1995). However, its low throughput and higher sample volume requirements impede its use for large screening campaigns. Fluorescence-based thermal shift methods provide a more accessible alternative for routine measurements, requiring smaller sample amounts and being well suited to medium- and high-throughput applications (Niesen et al., 2007; Kazlauskas et al., 2021; Gooran & Kopra, 2024). These methods can rely either on extrinsic fluorescent probes or on temperature-dependent changes in intrinsic protein fluorescence. Because terminology varies across the literature, with terms such as thermal shift assay (TSA), differential scanning fluorimetry (DSF) and Thermofluor sometimes used interchangeably (Petrauskas et al., 2024), we use here TSA for extrinsic-dye-based assays and DSF specifically for label-free measurements based on intrinsic protein fluorescence.

Commonly defined as the midpoint of the unfolding transition, *T*_m_ is widely used as a convenient descriptor of thermal stability. At equilibrium, for a fully reversible two-state transition, *T*_m_ corresponds to the temperature at which the folded and unfolded states are equally populated and the standard Gibbs free energy change of unfolding, Δ*G*°, is zero. Although *T*_m_ provides a simple and practical comparative metric, it does not by itself provide a complete thermodynamic description of protein stability (Niesen et al., 2007; Razban, 2019; Kotov et al., 2021). Experimentally determined transition temperatures may depend on conditions such as buffer composition, pH, heating rate and protein concentration (Niebling et al., 2021; Lisina et al., 2023). Their interpretation becomes particularly complex when unfolding is not fully reversible or is kinetically controlled (Niesen et al., 2007; Bai et al., 2019; Gao et al., 2020; Charvat et al., 2025). Under such conditions, experimentally derived *T*_m_ values are best regarded as apparent unless equilibrium and reversibility have been established (Niesen et al., 2007; Gao et al., 2020). In addition, the transition temperature assigned experimentally may depend on the signal used to monitor unfolding and on the analysis procedure used to determine the transition. This issue is particularly relevant for fluorescence-based thermal shift methods, in which protein unfolding can be monitored through different fluorescence signals.

In TSA, extrinsic probes such as SYPRO Orange or ANS can interact with hydrophobic surfaces exposed during thermal denaturation and may themselves influence the unfolding process (Hawe et al., 2008; Forneris et al., 2009; Gao et al., 2020; Gooran & Kopra, 2024). By contrast, the DSF approach considered here monitors heat-induced changes in the intrinsic fluorescence of aromatic residues. Tryptophan fluorescence is particularly sensitive to its local environment (Vivian & Callis, 2001): during thermal unfolding, red shift in the emission spectrum is often induced by exposure of buried tryptophan residues to solvent (Eftink, 1994), whereas blue shifts may reflect specific conformational rearrangements that place them in a less polar environment (Duy & Fitter, 2006). In proteins containing several tryptophan residues, the measured spectrum represents the combined contribution of residues experiencing different local environments. Tyrosine fluorescence can also report on conformational changes (Lee et al., 1992; Zhdanova et al., 2017), primarily through changes in intensity. However, its emission maximum is generally less sensitive to solvent polarity (Szabelski et al., 2002). As a result, thermal unfolding can produce complex changes in both fluorescence intensity and spectral shape.

In practice, these fluorescence changes can be represented using several readouts. The ratio of fluorescence intensities measured at 350 nm and 330 nm (F350/F330) is widely used, but alternative approaches include single-wavelength fluorescence intensities and full-spectrum approaches. The latter integrate scalar descriptors such as the barycentric mean (BCM) and multivariate methods like singular value decomposition (SVD). Because these readouts represent the fluorescence data in fundamentally different ways, they may not yield identical apparent transition parameters or provide equivalent information for quantitative analysis. Moreover, the suitability of a given fluorescence readout may depend on protein composition and on the spectral changes associated with the unfolding transition. The resulting thermal unfolding profiles can be analysed using first-derivative approaches or fitted with unfolding models to extract apparent transition parameters.

Beyond readout selection and data analysis, DSF measurements may also be influenced by instrument-specific factors, such as optical configuration, detector response, temperature calibration and thermal regulation. Similar concerns regarding inter-instrument comparability have been reported for other biophysical techniques (Ravi et al., 2010; Hole et al., 2013; Zhao et al., 2015; Velazquez-Campoy et al., 2021). However, the respective contributions of acquisition conditions, instrument-specific effects and analysis workflow to DSF variability remain insufficiently documented under interlaboratory conditions.

To address these questions, we conducted a two-phase interlaboratory DSF benchmarking study using a shared V_H_H sample across 26 laboratories and 28 instruments representing seven DSF instrument models. In Phase 1, participants used their routine acquisition and analysis workflows to capture variability associated with routine practice. In Phase 2, acquisition conditions were harmonised to reduce experimental heterogeneity and provide a more controlled basis for inter-instrument comparison. Results reported by participants were subsequently compared with values obtained after centralised reprocessing using MoltenProt (Burastero et al., 2021; Kotov et al., 2021) within a unified analysis workflow.

We assessed within-instrument repeatability and inter-instrument comparability of apparent thermal transition parameters and examined the respective contributions of acquisition conditions, fluorescence readout and data analysis. We further evaluated the robustness of apparent unfolding enthalpies derived from DSF and compared transition parameters obtained using different fluorescence readouts and analysis approaches with DSC measurements of the same V_H_H. Together, this design allowed us to examine both the reproducibility of DSF measurements across instruments and the extent to which the choice of fluorescence readout and analysis strategy influences the parameters extracted from thermal unfolding data.

## Results

### Phase 1: Global variability of DSF data across instruments

During this first interlaboratory phase, participating laboratories performed DSF measurements using their routine acquisition and analysis workflows. Apparent *T*_m_ values showed moderate inter-instrument variability with an overall mean of 58.3 ± 2.8 °C (**Figure 1a**; **Table 1)**. The corresponding pooled distribution showed that most *T*_m_ values clustered between 58 and 59 °C, despite a few more dispersed instrument-specific values (**Figure 1b**). To better describe the dataset distribution and reduce the influence of extreme values, median and interquartile range were also considered: the median *T*_m_ was 58.4 °C, with first quartile (Q1) and third quartile (Q3) at 57.6 °C and 59.2 °C, respectively (**Table 1**). Thus, the central 50% of measurements fell within a 1.6 °C interval, indicating a narrower core distribution than suggested by the overall standard deviation. Interestingly, repeated measurements acquired on the same physical instrument showed lower dispersion, with a median within-instrument interquartile range (IQR) of 0.2 °C (Q1–Q3, 0.1–0.4 °C). By contrast, apparent *T*_onset_ values reported by participants were more variable with a mean of 47.9 ± 6.2 °C (**Figure S2; Table 1**). Within-instrument dispersion was lower but remained substantial with a median IQR of 0.60 °C (Q1–Q3, 0.3–1.8 °C). Given this wider dispersion, *T*_onset_ was not considered in subsequent comparative analyses.

**Figure 1.**
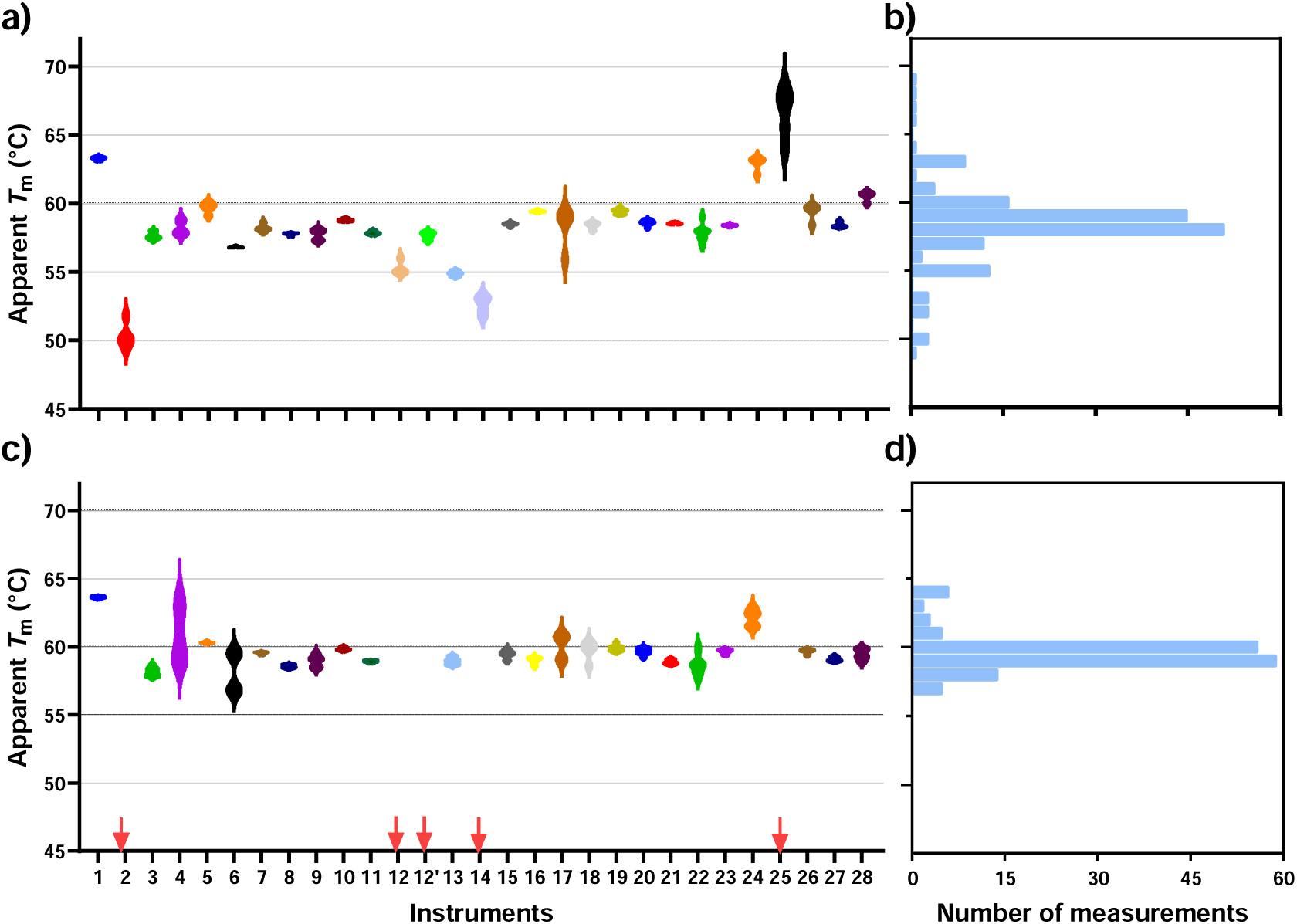
Apparent T_m_ distributions before and after MoltenProt reprocessing in Phase 1. Upper panels: Participant-reported apparent *T*_m_ values using their routine workflows, shown by instrument (a) and as a pooled distribution (b). Lower panels: Apparent *T*_m_ values obtained after centralised reprocessing with MoltenProt, shown by instrument (c) and as a pooled distribution of retained values (d). Violin plots show *T*_m_ distributions by instrument, whereas horizontal histograms show pooled distributions using a 1 °C bin width. Red arrows indicate instruments for which datasets were not retained after application of the MoltenProt quality control criteria. 12 and 12′ correspond to the same instrument but data were analysed using two different workflows.

**Table 1.** Summary statistics of participant-reported and MoltenProt-derived apparent transition temperatures in Phases 1 and 2. Values are reported as mean ± standard deviation (SD) and median [Q1–Q3]. Within-instrument interquartile range (IQR) values were calculated from replicate measurements for each instrument and are reported as median [Q1–Q3] across instruments.

| Parameter | Phase 1<br>mean ± SD | Phase 1 median<br>[Q1–Q3] | Phase 2<br>mean ± SD | Phase 2 median<br>[Q1–Q3] |
| --- | --- | --- | --- | --- |
| $T_m$ (°C) from participants | 58.3 ± 2.8 | 58.4 [57.6–59.2] | 58.0 ± 2.6 | 58.4 [57.1–59.5] |
| Median within-instrument<br>IQR of $T_m$ (°C) | – | 0.2 [0.10–0.4] | – | 0.2 [0.1–0.3] |
| $T_{onset}$ from participants | 47.9 ± 6.2 | 47.5 [44.4–51.5] | 47.1 ± 4.6 | – |
| Median within-instrument<br>IQR of $T_{onset}$ (°C) | – | 0.6 [0.3–1.8] | – | – |
| $T_m$ (°C) with MoltenProt | 59.5 ± 1.3 | 59.4 [58.8–59.8] | 59.5 ± 1.3 | 59.4 [59.0–59.8] |
| Median within-instrument<br>IQR of $T_m$ with MoltenProt<br>(°C) | – | 0.3 [0.2–0.5] | – | 0.2 [0.1–0.4] |

After centralised reprocessing, the apparent *T*_m_ distribution became narrower, with a mean of 59.5 ± 1.3 °C and a median of 59.4 °C [58.8–59.8] (**Figure 1c**; **Table 1**) for datasets that could be fitted with MoltenProt. This narrowing was also visible in the pooled distribution, which became more concentrated around 59–60 °C after reprocessing (**Figure 1d**). Thus, centralised reprocessing reduced the spread of apparent *T*_m_ values across retained datasets.

Overall, 145 out of 171 submitted datasets (85%) met MoltenProt internal quality criteria and were successfully fitted using the two-state unfolding model (**Table S1**). Datasets that were not retained generally showed low-intensity or poorly defined fluorescence transitions. Notably, datasets not retained after the centralised quality control procedure had nevertheless been assigned apparent *T*_m_ values by the participating laboratories, highlighting heterogeneity in local criteria for data interpretation and acceptance.

The relationship and agreement between participant-reported and MoltenProt-derived apparent *T*_m_ values were assessed by correlation and Bland–Altman analyses, respectively (**Figure 2**). The correlation plot showed an overall relationship between the two analyses (**Figure 2a**), while Bland–Altman analysis gave a mean bias of +1.0 °C with 95% limits of agreement from −1.3 °C to +3.3 °C. This indicates a small overall tendency towards higher apparent *T*_m_ values after MoltenProt reprocessing than in participant-reported analyses (**Figure 2b**). Overall, 91% of paired measurements fell within these limits of agreement.

**Figure 2.**
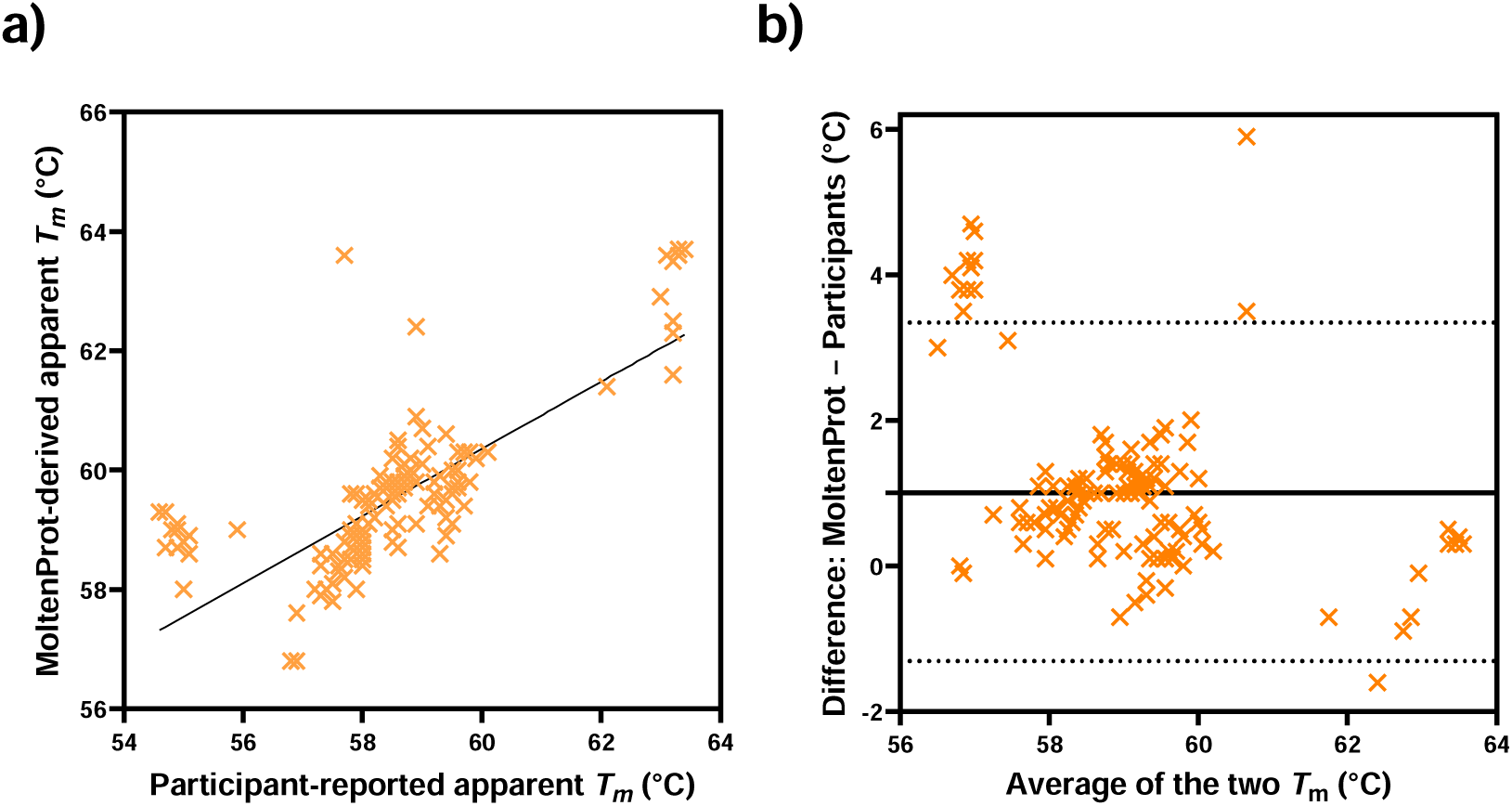
Relationship and agreement between participant-reported and MoltenProt-derived T_m_ across all Phase 1 datasets. (a) Correlation plot showing an overall relationship between participant-reported and MoltenProt-derived analyses. Black line: linear regression line. (b) Bland–Altman representation of the same data. The mean bias of ≈ +1 °C (solid line) and the 95% limits of agreement (dotted lines, –1.3 °C to +3.3 °C) indicate that MoltenProt yields slightly higher apparent *T*_m_ values than the participant analyses. 91% of the paired measurements fall within this interval.

Phase 2 was then initiated to explore the effect of harmonised experimental conditions on the variability observed.

### Phase 2: Comparison under harmonised acquisition conditions

In Phase 2, each laboratory used the same instrument as in Phase 1 but followed a harmonised acquisition protocol with a standardised temperature ramp and fluorescence settings adapted to each instrument. Under these harmonised conditions, participant-reported apparent *T*_m_ had a mean of 58.0 ± 2.6 °C with a median of 58.4 °C [57.1–59.5], remaining within the same range as those obtained in Phase 1 (**Figure 3a**; **Table 1**). The corresponding pooled distribution confirmed that most values again clustered around 58–60 °C, while a limited number of instrument-specific values remained more dispersed (**Figure 3b**). Replicate measurements on the same physical instrument showed similar dispersion to that observed in Phase 1, with a median within-instrument IQR of 0.2 °C (Q1–Q3, 0.1–0.3 °C).

**Figure 3.**
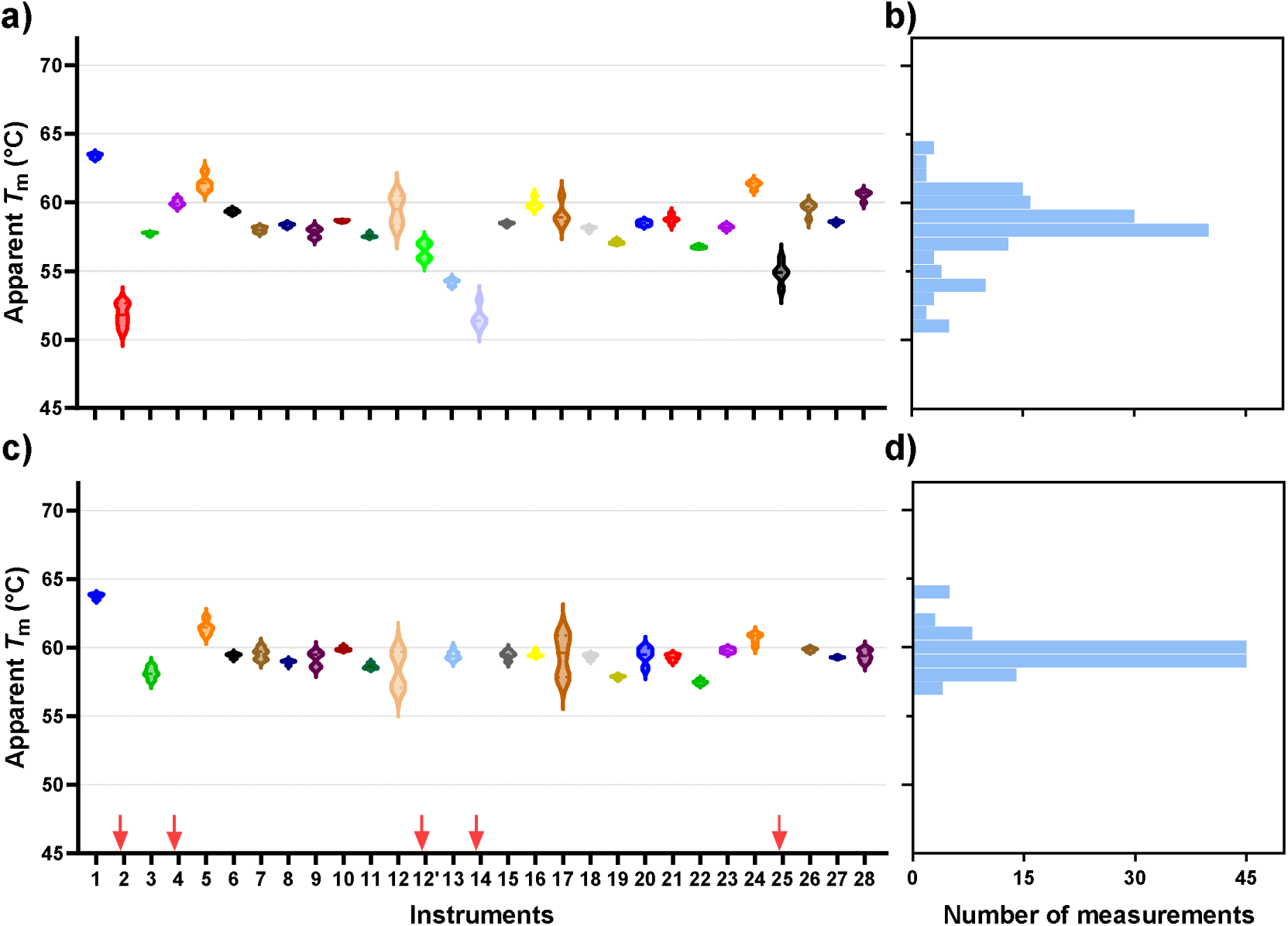
Apparent T_m_ distributions under harmonised acquisition conditions in Phase 2 before and after MoltenProt reprocessing. Upper panels: Apparent *T*_m_ values obtained from participants using their routine analysis workflows under harmonised acquisition conditions, shown by instrument (a) and as a pooled distribution (b). Lower panels: Apparent *T*_m_ values obtained after centralised reprocessing with MoltenProt, shown by instrument (c) and as a pooled distribution of retained values (d). Violin plots show *T*_m_ distributions by instrument, whereas horizontal histograms show pooled distributions using a 1 °C bin width. Red arrows indicate instruments for which datasets were not retained after application of the MoltenProt quality control criteria. 12 and 12′ correspond to the same instrument but data were analysed using two different workflows.

Centralised reprocessing with MoltenProt yielded a mean apparent *T*_m_ of 59.5 ± 1.3 °C with a median of 59.4 °C [59.0–59.8], again producing a narrower distribution among retained datasets than for participant-reported values and a pattern similar to that observed in Phase 1 (**Figure 3c**; **Table 1**). This effect was also visible in the pooled distribution, which became more concentrated around 59–60 °C after centralised reprocessing (**Figure 3d**). Within-instrument variability remained low after reprocessing, with a median within-instrument IQR of 0.2 °C (Q1–Q3, 0.1–0.4 °C). Overall dispersion across instruments remained similar to that observed in Phase 1, although differences persisted both between instruments of the same laboratory (**Table S2**) and between instruments of the same model. In summary, harmonising acquisition conditions did not drastically reduce the inter-instrument dispersion of apparent *T*_m_ values.

### DSF versus DSC: orthogonal comparison of thermal transitions

To provide an orthogonal calorimetric comparison, the same V_H_H sample was analysed by DSC. Fitting with a two-state model including a non-zero unfolding heat-capacity change (Δ*C*_p_) yielded a *T*_m_ of 55.5 ± 0.1 °C and an unfolding enthalpy of 95 ± 2 kcal/mol (**Figure 4a**). The heat capacity maximum occurred at 56.0 °C, slightly higher than the *T*m. For this V_H_H, the *T*_m_ measured by DSC was approximately 2.5–3 °C lower than the mean apparent *T*_m_ values reported by participants using DSF in Phases 1 and 2 (**Table 1**).

**Figure 4.**
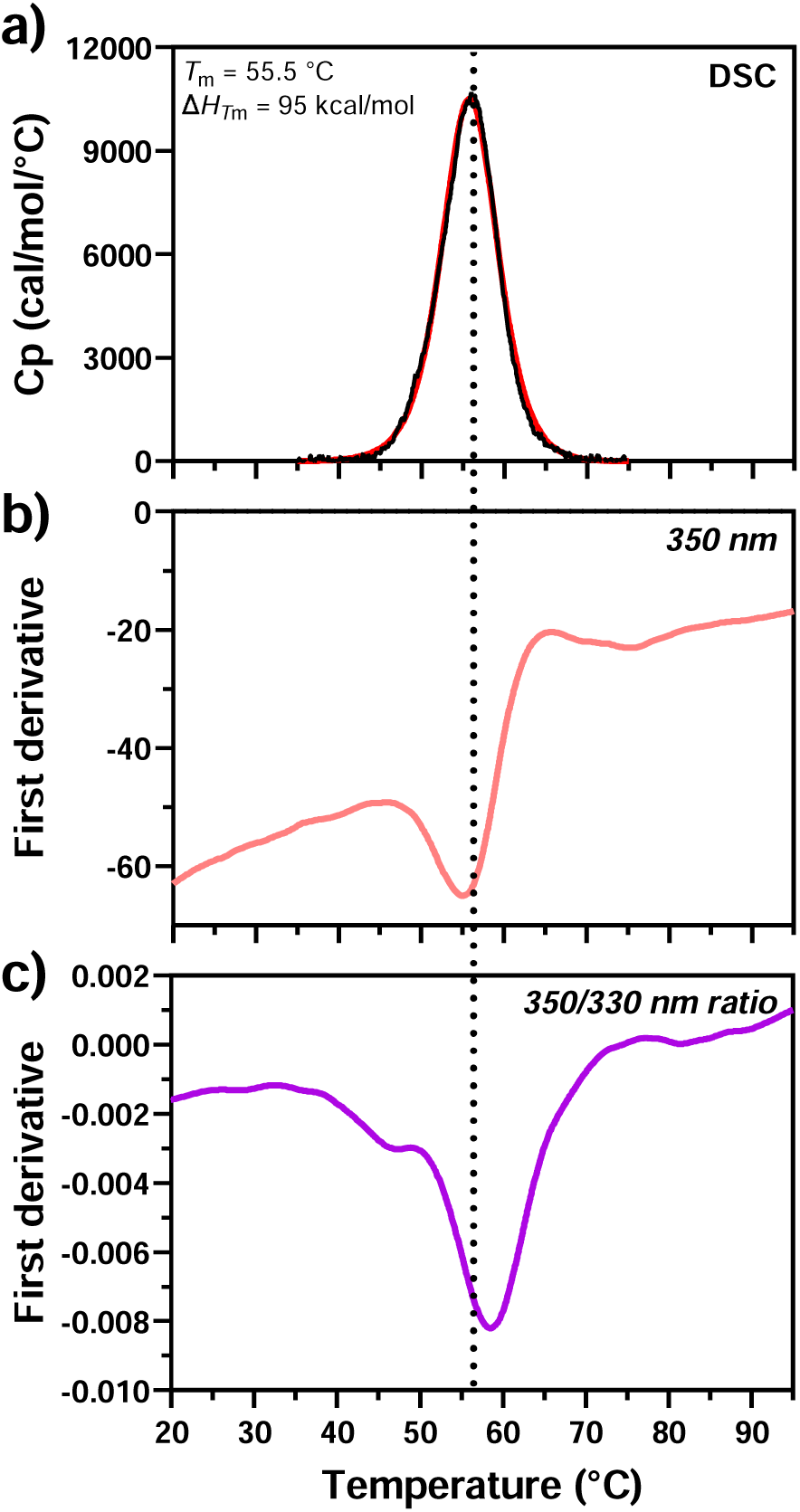
Comparison of the reference V_H_H thermal transition by DSC and DSF fluorescence readouts for instrument 15. DSF measurements were performed at 0.3 mg/mL and DSC at 0.4 mg/mL. (a) DSC thermogram showing a single cooperative transition, with a *T*_m_ of 55.5 °C and an unfolding enthalpy of 95 kcal/mol. (b) First-derivative trace of the DSF fluorescence signal recorded at 350 nm. The dashed vertical line indicates the *T*_m_ obtained by DSC. The apparent *T*m is close to this value. (c) First-derivative trace obtained from the 350/330 nm ratio under identical conditions. The apparent *T*_m_ is shifted towards a higher temperature.

When the DSC thermogram was examined alongside representative DSF traces for this V_H_H, the apparent *T*_m_ obtained from the fluorescence intensity at 350 nm (**Figure 4b**) was closer to the DSC transition temperature than the value obtained from the 350/330 nm ratio, which remained shifted to higher temperatures (**Figure 4c**). A similar pattern was also reflected in the interlaboratory dataset: datasets from instruments 12 and 13 which used the 350 nm fluorescence signal as their analysis readout reported apparent *T*_m_ values close to 55 °C, whereas datasets analysed using the 350/330 nm ratio or the BCM generally yielded higher apparent *T*_m_ values, typically ≥ 58 °C.

In addition, analysis of the same V_H_H sample on two different DSF instrument models within the same laboratory, under otherwise identical sample and acquisition conditions, produced apparent *T*_m_ differences of up to 7 °C, depending on the instrument, fluorescence readout and fitting method (**Table S2**). Together, these observations illustrate, for this V_H_H, how the apparent agreement between DSF and DSC depends strongly on the fluorescence readout and analysis strategy.

### Apparent unfolding enthalpies derived from DSF

To assess whether DSF datasets could also provide apparent quantitative information beyond apparent *T*_m_, apparent enthalpies of unfolding were extracted from the 350/330 nm ratio with MoltenProt websever, using an equilibrium two-state model. When fitting was restricted to a temperature window centred on the unfolding transition (40–80 °C), the mean apparent Δ*H* across the retained DSF datasets was 87.5 ± 19.3 kcal/mol and the median was 84.4 kcal/mol [78.8–92.2] (**Figure 5**; **Table 2**). For comparison, DSC analysis yielded an unfolding enthalpy of 95 kcal/mol. The pooled distribution showed substantial heterogeneity of apparent Δ*H*, with most values falling around 70–100 kcal/mol and a tail extending towards higher values (**Figure 5b**). The values were in the range of the DSC value, but with substantial heterogeneity across instruments and datasets. However, the median within-instrument IQR was 4.4 kcal/mol [1.9–10.2], indicating variability among replicate measurements. To examine the influence of fluorescence readouts, instrument 28 was selected because its full spectral acquisition mode allowed the same measurements to be analysed using single-wavelength fluorescence at 350 nm, the 350/330 nm ratio, BCM and SVD. Within the same 40–80 °C fitting window, the estimated apparent enthalpy differed by up to 20% depending on the fluorescence readout (**Table S3**). As a sensitivity analysis, the instrument 28 datasets were additionally fitted over the full acquisition range (20–95 °C). Changing the fitting interval alone altered the estimated apparent enthalpy by up to 30% (**Table S3**).

**Figure 5.**
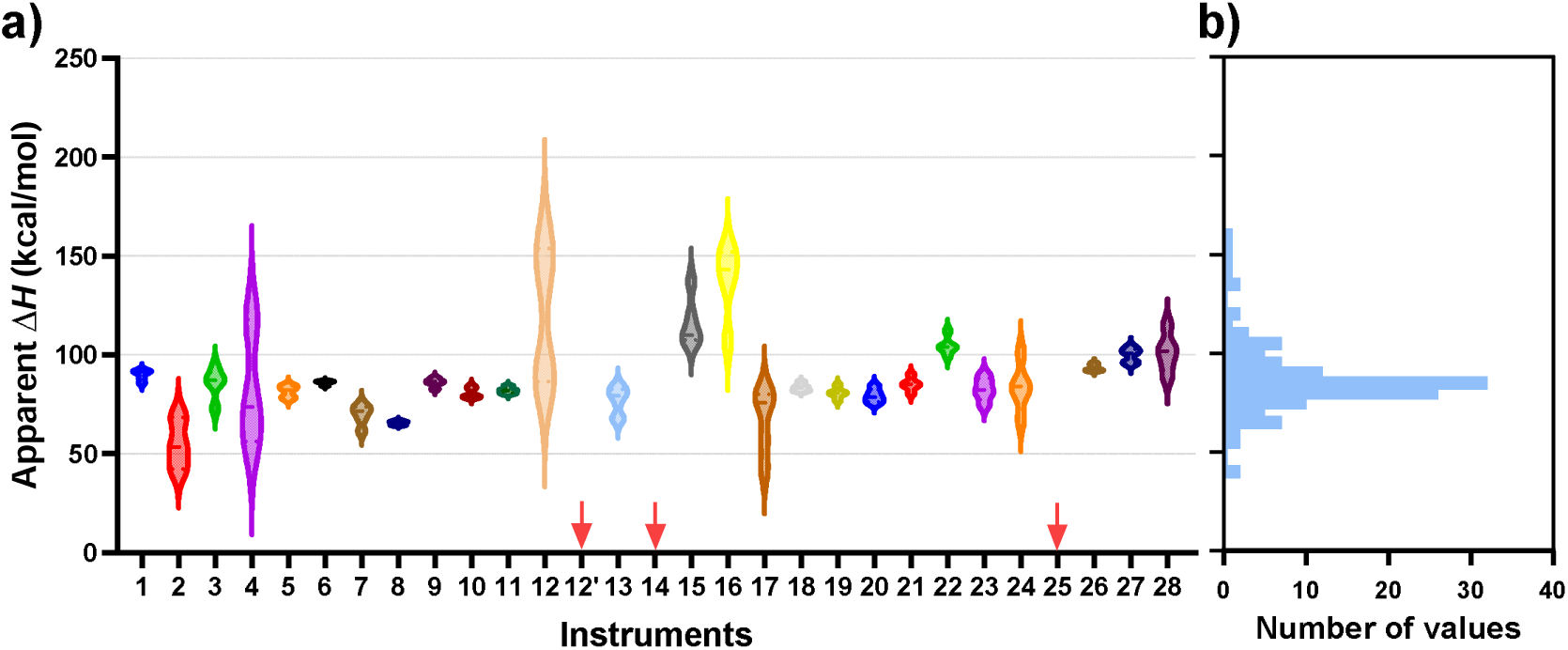
Apparent ΔH values across instruments using the 40–80 °C fitting range. Apparent Δ*H* values were derived from the 350/330 nm fluorescence ratio by fitting the thermal unfolding profiles with the two-state equilibrium model implemented in MoltenProt over the 40–80 °C temperature range. (a) Distribution of apparent Δ*H* values for each instrument. Each violin plot represents replicate measurements obtained for a given instrument. (b) Pooled distribution of individual apparent Δ*H* values across all retained datasets, shown using a 10 kcal/mol bin width. The overall mean apparent Δ*H* was 87.5 ± 19.3 kcal/mol. Red arrows indicate instruments for which datasets were not retained after application of the MoltenProt quality control criteria.

**Table 2.**
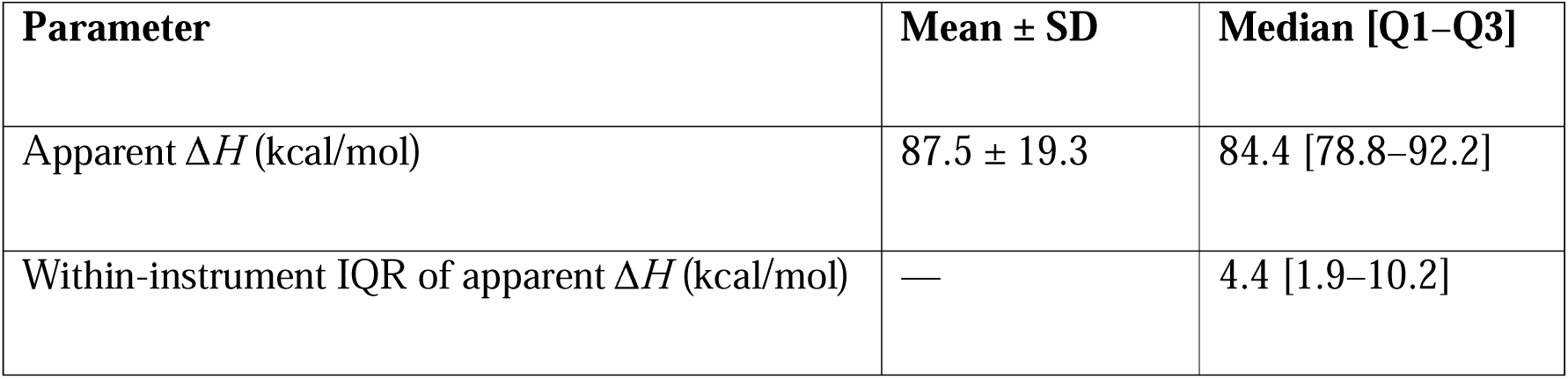
Summary statistics of DSF-derived apparent ΔH values across retained datasets. Apparent Δ*H* values were derived from the 350/330 nm fluorescence ratio by fitting the thermal unfolding profiles with the two-state equilibrium model implemented in MoltenProt over the 40–80 °C temperature range. Mean ± SD and median [Q1–Q3] were calculated across individual retained datasets. Within-instrument IQR values were calculated from replicate measurements for each instrument and are reported as median [Q1–Q3] across instruments. For comparison, DSC analysis of the same V_H_H yielded an unfolding enthalpy of 95 kcal/mol.

| Parameter | Mean ± SD | Median [Q1–Q3] |
| --- | --- | --- |
| Apparent $\Delta H$ (kcal/mol) | 87.5 ± 19.3 | 84.4 [78.8–92.2] |
| Within-instrument IQR of apparent $\Delta H$ (kcal/mol) | — | 4.4 [1.9–10.2] |

Overall, DSF-derived apparent enthalpies showed substantial variability across the interlaboratory dataset. The extended analysis of instrument 28 further demonstrated the influence of fluorescence readout and fitting interval on the estimated values.

## Discussion

This collaborative interlaboratory benchmark was designed to assess how reproducible DSF measurements are in routine practice, at the level of individual instruments and across instruments and laboratories. Furthermore, we examined how fluorescence readout and analysis conditions influence the parameters extracted from DSF measurements. By combining a two-phase study design, centralised reprocessing in MoltenProt (Kotov et al., 2021), and an orthogonal calorimetric comparison with DSC, the study highlights three main findings. First, apparent *T*_m_ values are highly repeatable within a given instrument but are not fully transferable across instruments. Second, part of the observed variability arises from differences in data interpretation, fluorescence readout selection, and analysis practice rather than from acquisition conditions alone. Third, apparent enthalpies derived from DSF are much less robust than apparent *T*_m_ and should therefore be interpreted with greater caution.

### 1. DSF-derived apparent *T*_m_ is repeatable within a given instrument but not fully transferable across instruments

Across 26 laboratories and seven DSF instrument models, apparent *T*_m_ values for the reference V_H_H showed moderate dispersion, with mean values of 58.3 ± 2.8 °C and 58.0 ± 2.6 °C in Phases 1 and 2, respectively. Distribution-based analysis further indicated that the central 50% of values fell within relatively narrow ranges, with medians of 58.4 °C [57.6–59.2] and 58.4 °C [57.1–59.5] in Phases 1 and 2, respectively. Variability was slightly greater in Phase 2, in which all participants followed a common protocol. This suggests that the variability does not arise primarily from differences in experimental protocols but rather from instrument-specific effects, analysis differences or both. At the same time, replicates performed on the same instrument showed much lower dispersion, with a median within-instrument IQR of approximately 0.2 °C in both phases. Together, these results indicate high within-instrument repeatability but more limited quantitative comparability across instruments and laboratoraties.

Apparent *T*_m_ shifts of only 1–2 °C may be informative in formulation studies, ligand screening, or comparative assessments of protein stability within a controlled workflow. However, such small differences should be interpreted with caution when values are compared across instruments or with the literature.

Interestingly, centralised reprocessing with MoltenProt narrowed the apparent *T*_m_ distribution in both phases. This indicates that part of the interlaboratory variability came from differences in local transition assignment, fitting procedures and data-acceptance criteria. This interpretation is reinforced by the observation that some datasets excluded during centralised reprocessing because of weak or noisy traces had nevertheless been assigned apparent *T*_m_ values by participating laboratories. In other words, part of the variability did not arise from the measurement itself, but from the decision to extract a *T*_m_ from traces that did not support such an assignment.

At the same time, harmonising acquisition conditions in Phase 2 did not substantially reduce the overall inter-instrument dispersion relative to Phase 1. This suggests that interlaboratory variability was driven by both instrument-specific and analysis-dependent factors. Differences remained between instruments within the same laboratory and between instruments of the same model, even when the same fluorescence readout was used. Factors such as optical geometry, detector response, temperature calibration, and temperature control are therefore likely to contribute to the variability. An additional explanation for the limited effect of Phase 2 harmonisation is that many laboratories were already operating under broadly similar manufacturer-recommended conditions, such that a degree of practical similarity already existed in routine use. Overall, protocol harmonisation alone is therefore not sufficient to ensure quantitative comparability across instruments.

Improved comparability will require better control of instrument-specific offsets and improved training in DSF data interpretation, including explicit quality-control criteria. As in mass spectrometry or circular dichroism, where reference materials are routinely used for calibration, DSF would benefit from a well-characterised protein standard with a consensus *T*_m_, analysed alongside unknown samples to correct for possible instrument-specific offsets. Until such standards are established, DSF-derived apparent *T*_m_ values should primarily be used comparatively within a defined instrument and analysis workflow.

### 2. Comparison with DSC illustrates the dependence of DSF-derived apparent *T*_m_ on fluorescence readout and analysis mode

A central finding of the study emerges from the comparison with DSC. For this V_H_H, DSC displayed a single cooperative transition with a *T*_m_ of 55.5 °C, whereas the mean apparent *T*_m_ across DSF instruments was approximately 58 °C. Thus, under the conditions examined here, apparent *T*_m_ extracted from DSF measurements did not necessarily coincide with the transition temperature obtained by DSC. Such differences are not unexpected, since the temperature assigned to a thermal transition can depend on the experimental observable and on the criterion used to assign the transition temperature. In particular, the temperature of maximal excess heat capacity measured by DSC may not coincide exactly with the inflection temperature of a spectroscopic unfolding curve, and their relationship depends on the thermodynamic parameters of the transition, including the unfolding enthalpy and heat-capacity change (Velazquez-Campoy et al., 2016). The DSC comparison should therefore be regarded here as an orthogonal reference for examining how DSF-derived transition parameters depend on fluorescence readout and analysis strategy.

Notably, the offset relative to DSC was not uniform across fluorescence readouts and analysis modes. For this V_H_H, single-wavelength analysis at 350 nm produced apparent *T*_m_ values closer to the transition temperature observed by DSC, whereas ratio and BCM approaches generally produced higher values. This observation identifies 350 nm as an informative readout for the present sample but should not be interpreted as evidence that this wavelength is universally optimal for DSF analysis. In routine practice, participating laboratories used a range of fluorescence readouts, often reflecting instrument defaults or established local workflows. Ratio-based analyses were particularly common, but for the V_H_H examined here they did not generally provide the closest agreement with the DSC transition. These findings therefore highlight the importance of evaluating the suitability of the fluorescence readout for the protein and experimental context under study.

These observations have a mechanistic basis. A fundamental requirement for quantitative analysis of an equilibrium unfolding transition is that the experimental observable be related in a defined manner to the populations of the underlying macrostates. Fluorescence intensity measured at a fixed emission wavelength provides the most straightforward example, since the observed signal can be expressed as the population-weighted contribution of the folded and unfolded states. By contrast, fluorescence intensity ratios and spectrum-derived quantities such as BCM are nonlinear combinations of the underlying fluorescence signals and therefore do not in general preserve a linear relationship with state populations. Consequently, although such parameters may provide useful empirical descriptors of a transition and can make a conformational transition easier to detect, quantitative parameters derived from them should be interpreted with caution. This is consistent with previous theoretical and experimental work questioning the use of fluorescence ratios and other derived fluorescence observables for quantitative analysis of protein unfolding transitions (Eftink, 1994; Žoldák et al., 2017).

Importantly, however, the suitability of any fluorescence observable remains protein- and context-dependent. For proteins lacking tryptophan, for example, large environment-sensitive spectral changes may be absent, limiting the suitability of some intrinsic-fluorescence readouts (Szabelski et al., 2002). Fluorescence responses may also be affected by changes in spectral shape, wavelength-dependent quenching or enhancement, or by ligands and formulation components that influence different spectral regions unequally. Under such conditions, the 350/330 nm ratio reflects not only redistribution between folded and unfolded populations but also wavelength-dependent changes in intrinsic fluorescence. Conversely, in the absence of a significant spectral shift, the ratio may remain nearly constant even when a clear unfolding transition is visible at a single wavelength.

Full-spectrum approaches require a similar distinction. BCM can provide a useful description of global spectral displacement but, being a derived scalar parameter, it does not necessarily track state populations linearly and therefore cannot automatically be assumed to yield quantitative parameters. By contrast, multivariate approaches such as SVD retain more of the structure of the spectral dataset and can reveal coherent spectral changes that may be obscured by simple ratios or scalar descriptors (Gray et al., 2012; Kunka et al., 2022). In the extended analysis of instrument 28, SVD-derived apparent *T*_m_ values did not show the upward shift observed with the BCM and the 350/330 ratio and were similar to those obtained from single-wavelength fluorescence at 350 nm. Thus, for this V_H_H, both SVD and 350 nm yielded apparent *T*_m_ values closer to the DSC transition temperature than ratio- or BCM-based analyses. For apparent Δ*H*, a different pattern emerged: estimates derived from SVD and single-wavelength fluorescence at 350 nm were less sensitive to the choice of fitting range than ratio- or BCM-based estimates (**Table S3**). This lower sensitivity to the fitting range should not, however, be interpreted as evidence of greater thermodynamic accuracy. Although these observations are limited to the present sample, they suggest that SVD represents a promising full-spectrum approach that warrants further evaluation across proteins with different intrinsic fluorescence properties and under different experimental conditions.

Thus, the present benchmark does not imply that ratio or BCM analyses are inappropriate for routine DSF screening. Rather, it highlights the need to distinguish between fluorescence readouts that are useful for detecting or comparing thermal transitions and those suitable for quantitative interpretation of the transition parameters. In this respect, our results reinforce previous concerns regarding the use of fluorescence ratios and other derived spectral parameters without appropriate validation and underline the importance of selecting an observable with a well-defined relationship to state populations.

### 3. Apparent enthalpies derived from DSF are less robust than apparent *T*_m_

Beyond *T*_m_, the benchmark also examined whether DSF could provide informative estimates of apparent unfolding enthalpy. When fitting was restricted to the unfolding region (40– 80 °C), the mean Δ*H* derived from DSF was 87.5 ±19.3 kcal/mol and numerically close to the DSC value of 95 ±19.3 kcal/mol for this V_H_H. However, the distribution was broad and far more dispersed than that observed for apparent *T*_m_. This indicates limited robustness across instruments and datasets.

Apparent Δ*H* derived from DSF also depended strongly on the fluorescence readout and fitting window. In the extended analysis of instrument 28, changing the fitting range from 40– 80 °C to 20–95 °C altered apparent Δ*H* substantially for some readouts (**Table S3**). This outcome highlighs the sensitivity of these estimates to the pre- and post-transition regions used to constrain the fit.

These estimates were obtained using the two-state equilibrium model implemented in MoltenProt, which was selected because it calculates an apparent unfolding enthalpy, unlike the empirical two-state model, which uses *T*_onset_ as a descriptive parameter. This distinction is particularly relevant here given the much broader variability observed for *T*_onset_ across the benchmark dataset.

This sensitivity is expected from a thermodynamic perspective. Unlike DSC, which directly measures heat-capacity changes and therefore provides thermodynamic observables directly related to the unfolding process (Durowoju et al., 2017), DSF reports optical consequences of conformational changes. In practice, DSF-derived apparent enthalpies may therefore remain useful for internal ranking or relative comparison within a tightly controlled workflow, but the present results indicate that such values should be treated with considerable caution in inter-instrument or cross-study comparisons.

### 4. Limitations of the present benchmark

Several limitations should be acknowledged. First, the study was restricted to one model protein, a V_H_H displaying a single cooperative unfolding transition. While this facilitated interlaboratory comparison, it limits the generalisation of the conclusions to more complex systems. Second, instrument models were not equally represented across the benchmark. Consequently, pooled statistics are weighted towards the most represented models and should not be interpreted as a balanced comparison across DSF instrument models. Third, the comparison with DSC was based on a single calorimetric dataset for this sample. It should therefore be regarded as an orthogonal comparison illustrating how parameters derived from DSF measurements depend on the fluorescence readout and analysis strategy, rather than as a general assessment of systematic agreement or bias between DSF and DSC. Finally, DSF measurements are performed during continuous heating and therefore do not necessarily probe equilibrium unfolding. This is particularly important because heat-induced unfolding is frequently irreversible, in which case equilibrium thermodynamic parameters cannot be rigorously extracted from the transition. Moreover, even for a reversible transition, equilibrium will only be maintained if the heating is sufficiently slow relative to the kinetics of conformational equilibration, which are intrinsically protein-dependent. Consequently, scan-rate dependence should be considered when interpreting DSF data, and *T*_m_ and especially Δ*H* derived under non-equilibrium conditions should be regarded as apparent parameters.

### 5. Practical recommendations for DSF data reporting and comparison

The present benchmark highlights several practical points that may improve the comparability of DSF datasets across laboratories. First, DSF studies should report, as explicitly as possible, the fluorescence readout used for analysis, the fitting model, the temperature interval used for analysis and the criteria applied for data acceptance or rejection.

Second, apparent *T*_m_ differences on the order of 1–2 °C or less should therefore be interpreted with caution when data are compared across instruments or laboratories. Indeed, differences of this magnitude may in some cases reflect instrument- or analysis-dependent offsets rather than real differences.

Third, the use of a well-characterised reference material that could be widely adopted across DSF users would provide a practical means of monitoring instrument performance and detecting systematic temperature offsets or instrumental drift. Such an approach could support validated recalibration procedures and should be combined with transparent reporting of acquisition conditions, fluorescence readout, analysis methods and data-acceptance criteria to improve inter-instrument comparability of DSF measurements.

## Conclusion

This interlaboratory benchmark shows that DSF measurements can be highly repeatable within a given instrument and workflow, but that apparent *T*_m_ values are not fully transferable across instruments or laboratories. The residual variability observed after harmonisation indicates contributions from both analysis-dependent and instrument-specific factors. For the V_H_H studied here, agreement with DSC depended strongly on the fluorescence readout and analysis strategy, while DSF-derived apparent unfolding enthalpies were markedly less robust than apparent *T*_m_. These results support the development and use of shared reference materials for routine monitoring of instrument performance, together with transparent reporting of acquisition and analysis settings to improve inter-instrument comparability of DSF measurements.

## Materials and methods

### Participating laboratories and instruments

Twenty-six partner laboratories participated in the study, using a total of 28 instruments representing seven DSF instrument models (**Figure S1**). The instrument distribution was as follows: Prometheus NT.48 (NanoTemper Technologies GmbH, n = 11), Prometheus Panta (NanoTemper Technologies GmbH, n = 7), SUPR-DSF (Nicoya, n = 4), Uncle (Unchained Labs, n = 3), Cary Eclipse (Agilent Technologies, n = 1), Aunty (Unchained Labs, n = 1) and Prometheus NT.Plex (NanoTemper Technologies GmbH, n = 1).

### Common reference sample

A common V_H_H reference sample was used throughout the benchmarking study to minimise variability between participants (Birchenough et al., 2021). This anti-lysozyme V_H_H was developed through the European MOSBRI project prior to distribution to participating sites. Before shipment, the quality of the lyophilised V_H_H was checked to verify its homogeneity, purity, and integrity. In brief, dynamic light scattering was used to assess sample homogeneity, UV absorbance spectroscopy to determine protein concentration and evaluate possible nucleic acid contamination, and MALDI-TOF mass spectrometry to confirm molecular integrity and purity. This quality control characterisation package was adapted from previously published approaches (Raynal et al., 2014; de Marco et al., 2021). Detailed methodological considerations are beyond the scope of the present study.

### Interlaboratory DSF benchmark workflow

The interlaboratory benchmark was conducted according to a two-step protocol distributed to all participating laboratories. This protocol, provided as **Supplementary Protocol S1**, describes the overall benchmarking strategy, the V_H_H sample preparation procedure, the acquisition parameters for both the user-defined and harmonised phases, and the results obtained.

#### a) Sample preparation

Participants received a pre-characterised lyophilised V_H_H sample together with a product information sheet describing the reconstitution procedure in water. Each laboratory was instructed to determine the concentration of the reconstituted V_H_H at 280 nm by recording an absorbance spectrum between 240 and 340 nm using a standard 1 cm pathlength cuvette. The V_H_H sample was then diluted in PBS to a final concentration of 0.3 mg/mL and the final concentration was verified by UV-visible spectroscopy before DSF acquisitions.

#### b) Phase 1: User-defined DSF protocols

In Phase 1, participants were asked to perform five replicate thermal unfolding measurements using their routine DSF acquisition settings and preferred analysis workflow. Laboratories reported the main acquisition and analysis parameters (e.g., instrument manufacturer and model, software used, excitation/emission settings, temperature range, ramp rate, equilibration time) together with the apparent *T*_m_ and onset temperatures (*T*_onset_). In this study, *T*_m_ values correspond to the transition temperatures assigned locally by each laboratory, typically from a derivative extremum, a maximal-slope temperature, or a fitted transition midpoint, depending on the instrument software and local workflow. *T*_onset_ corresponds to the locally assigned temperature at which the fluorescence change associated with unfolding was considered to start.

No restrictions were imposed on the temperature gradient, excitation source, fluorescence settings or analysis software, to reflect routine laboratory practice as closely as possible.

#### c) Phase 2: Harmonised DSF protocol

In Phase 2, all participating laboratories followed a common acquisition protocol. Each laboratory performed five replicate measurements using the V_H_H sample adjusted to 0.3 mg/mL. Measurements were performed using a linear temperature ramp of 1 °C/min over a 20–95 °C temperature range, with an equilibration time of 5 min before the start of the thermal scan. Where allowed by the instrument configuration, fluorescence was recorded using excitation at 280 nm and emission between 300 and 400 nm, or the closest equivalent setting provided by the manufacturer. Participants were also instructed to minimise bubble formation during loading and to allow sufficient lamp stabilisation before acquisition.

#### d) Data collection and processing

For both phases, participants submitted raw and processed fluorescence data, absorbance spectra recorded before and after dilution, and a brief description of the algorithm or software used to determine apparent *T*_m_ values. The reporting template provided to participants is included in **Supplementary Protocol S1**.

#### e) Centralised data processing

Upon receipt, all fluorescence datasets were reprocessed centrally at Institut Pasteur using MoltenProt, available at the eSPC platform (spc.embl-hamburg.de). For the centralised comparison of apparent *T*_m_ values, the same fluorescence readout as that reported by the participating laboratory was retained and analysed using a unified workflow. This approach avoided introducing a change in fluorescence readout when assessing the effect of a common analysis procedure. Thermal unfolding profiles were fitted over the 40–80 °C temperature range using the two-state equilibrium model implemented in MoltenProt.

For the interlaboratory analysis of apparent unfolding enthalpy, apparent Δ*H* values reported in Figure 5 and Table 2 were derived specifically from the 350/330 nm fluorescence ratio using the same two-state model and 40–80 °C fitting interval. In this framework, the fluorescence signal is modelled assuming an apparent equilibrium between folded and unfolded states, together with temperature-dependent baselines for the pre- and post-transition regions. This fitting procedure was used to estimate apparent *T*_m_ and apparent Δ*H* values. In this study, derived apparent enthalpy refers to the enthalpy parameter returned by the MoltenProt two-state fitting model when applied to fluorescence unfolding traces. Because unfolding reversibility was not assessed and the DSF model used here did not include an unfolding heat-capacity change, these values should be interpreted as apparent van’t Hoff-like enthalpies rather than as calorimetric unfolding enthalpies. MoltenProt automatic post-fitting quality control criteria were applied to determine which datasets were not retained for subsequent analyses. In these cases, MoltenProt was not able to fit the data with fit parameters within predefined bounds (25 < *T*_m_ < 85 °C and 20 < Δ*H* < 450 kcal/mol). Instrument 28 was then selected for additional readout-dependent analyses because its full spectral dataset allowed the same measurements to be analysed using multiple fluorescence readouts, including single-wavelength intensity at 350 nm, the 350/330 nm ratio, BCM and SVD. This comparative analysis across multiple readouts was not possible for instruments recording only a limited number of fluorescence channels.

#### f) Statistical analysis

For each dataset, the fitting results obtained with MoltenProt were compiled and compared to the values initially reported by participating laboratories. Statistical analyses were conducted using GraphPad Prism version 10. Agreement between participant-reported *T*_m_ values and MoltenProt-derived *T*_m_ values was assessed using two complementary approaches: linear correlation analysis to evaluate the overall relationship between the two datasets and Bland– Altman analysis to quantify the mean bias and 95% limits of agreement between the two methods.

### DSC measurements

The thermal unfolding of the V_H_H was assessed by differential scanning calorimetry to provide an orthogonal calorimetric comparison with the parameters derived from DSF. The molar excess heat capacity of the protein was measured as a function of temperature in an Auto-PEAQ-DSC differential scanning calorimeter (MicroCal, Malvern-Panalytical) with a sample at 29 μM (0.4 mg/mL) and a temperature scanning rate of 1 °C/min. Buffer scans were performed beforehand until scan reproducibility was observed. After concentration normalisation and baseline subtraction, the data were analysed using a two-state unfolding model allowing a non-zero unfolding heat-capacity change (Δ*C*_p_). This fit was used to estimate the unfolding temperature (*T*_m_) and the unfolding enthalpy (Δ*H*).

## Broader audience statement

Differential scanning fluorimetry (DSF) is widely used to compare protein stability, but results are often compared across laboratories and instruments. This interlaboratory benchmark shows that DSF is repeatable within a controlled workflow, but that inter-instrument comparisons require caution. The study highlights the need for reference materials, instrument-performance monitoring and transparent reporting to improve the reproducibility of DSF measurements.

## Abbreviations and symbols

DSF, TSA, DSC, V_H_H, *T*_m_, *T*_onset_, Δ*H*, Δ*C*_p_, BCM, SVD, IQR, SD, QC.

## Supplementary material description

This section includes the protocol distributed to participating laboratories for the MOSBRI interlaboratory DSF benchmarking study (**Supplementary Protocol S1**), the distribution of DSF instruments across participating laboratories (**Figure S1**), the participant-reported *T*_onset_ values in Phase 1 (**Figure S2**), the proportion of datasets retained after MoltenProt quality-control fitting (**Table S1**), the variation in apparent *T*_m_ values across DSF instruments and analysis methods (**Table S2**), and the apparent Δ*H* values obtained from different fluorescence readouts and fitting intervals (**Table S3**).

## Acknowledgements

The authors thank MOSBRI for supporting the organisation of this collaborative benchmarking study. This study has received funding from the European Union’s Horizon 2020 research and innovation programme under grant agreement No 101004806 (MOSBRI project). We acknowledge the Robotein^®^ platform of the BE Instruct-ERIC Centre for providing access to their equipment. The work at ULiège was supported by the Walloon Region and the European Regional Development Fund (ERDF) [Grant portfolio PHENIX n°244, project PHENIX_Analytique_ULiège n°397]. Measurement at CF Biomolecular Interactions and Crystallography of CIISB, Instruct-CZ Centre, was supported by MEYS CR (LM2023042) and European Regional Development Fund-Project “Innovation of Czech Infrastructure for Integrative Structural Biology” (No. CZ.02.01.01/00/23_015/0008175). This project has received funding from the European Union’s Horizon 2020 research and innovation programme under the Marie Skłodowska-Curie grant agreement No 945405. The authors also thank Giuliana Sarno and Ida Pacilio from Università di Salerno for their advice.

## Author contributions

Conceptualisation: S.B., M.G.-A., B.R., A.M.,

Methodology: S.B., M.C., B.R.

Data curation: S.B.

Formal analysis: S.B., S.V., A.V.-C.

Software: M.G.-A., O.B.

Investigation: S.B., M.C., A.S.G., O.A., S.V., A.V.-C., G.A., A.F., N.Ž.K., M.B., D.M., H.B.,

T.J., A.K.B., T.C., A.C., N.D., S.P., M.E., E.F., E.G.P., J.H., E.P., A.K., A.L., W.S., K.Lap.,

K.L., A.L.R., B.L.-M., R.M., C.M., M.V., A.H.M., R.W.W., G.S., P.O.T., S.U., M.A.-N., R.G.

Resources: All authors

Project administration: S.B., M.C., B.R.

Supervision: B.R.

Funding acquisition: P.E.

Visualisation: S.B., M.C., B.R.

Writing – original draft: S.B., M.C., B.R.

Writing – review & editing: All authors.

## Manuscript information

38 main-manuscript pages; 5 figures; 2 tables. Supplementary material: 9 pages, including 2 supplementary figures, 3 supplementary tables, and Supplementary Protocol S1.

## Supplementary material

DSF_benchmarking_Protein_Science_Supplementary_Material.pdf, containing Figures S1– S2, Tables S1–S3 and Supplementary Protocol S1.

## Data availability statement

Participant DSF datasets, experimental metadata and reprocessed files underlying the figures and tables are available by contacting the corresponding authors.

