## Supplementary material for "Interlaboratory benchmarking of differential scanning fluorimetry reveals sources of variability in protein thermal stability parameters": Supp info

**Index of supplementary information:**

**Supplementary figures:**

**Figure S1:** *Distribution of 28 DSF instruments across the 26 participating laboratories.*

**Figure S2:** *T_onset_ (DSF) values reported by participants in Phase 1.*

**Supplementary tables:**

**Table S1:** *Proportion of submitted datasets meeting MoltenProt quality‑control and convergence criteria after centralised reprocessing.*

**Table S2:** *Effect of fluorescence readout and analysis method on apparent T_m_ for two DSF instruments measured in the same laboratory.*

**Table S3:** *Effect of fluorescence readout and fitting range on apparent ΔH for instrument 28.* Values are mean ± SD (kcal/mol).

**Supplementary Protocol**

**Supplementary figures:**

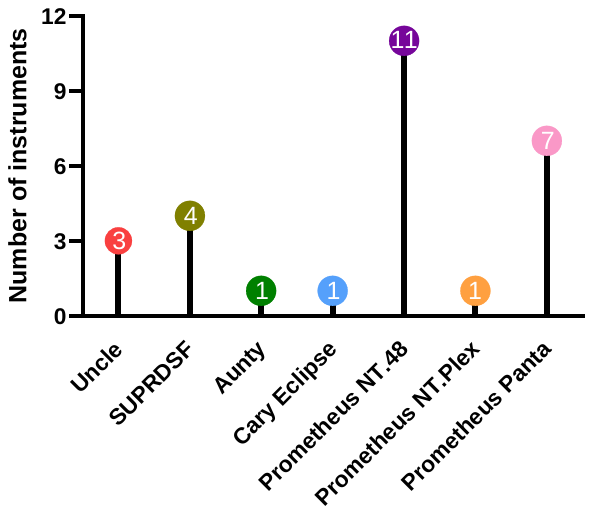

**Figure S1.** *Distribution of 28 DSF instruments across the 26 participating laboratories.*

Each coloured lollipop represents one instrument model, with the number inside each circle indicating the number of instruments of that model included in the benchmark.

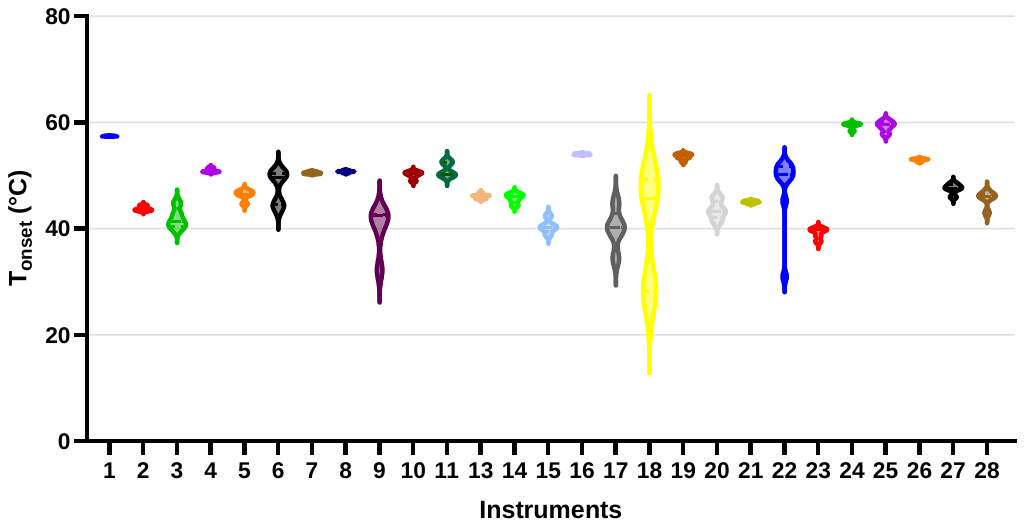

**Figure S2.** *T_onset_ (DSF) values reported by participants in Phase 1.*

Violin plots show the distribution of *T*_onset_ temperatures (°C) obtained by each instrument used across participating laboratories. Each colour represents a different DSF instrument model. The broad distribution observed for several instruments illustrates the greater dispersion of participant‑reported *T*_onset_ values. No *T*_onset_ values were reported for instrument 12.

**Supplementary tables:**

**Table S1.** *Proportion of submitted datasets meeting MoltenProt quality‑control and convergence criteria after centralised reprocessing.* “Datasets retained (%)” indicates the fraction of submitted datasets meeting the internal quality‑control (QC) and convergence criteria of the MoltenProt fitting workflow under the reprocessing conditions used in this study. Participants were requested to provide five replicate measurements; however, the number of submitted datasets varied among laboratories, with most laboratories providing five measurements and some providing up to ten replicates in total. These values reflect dataset compatibility with the analysis workflow applied here and should not be interpreted as an absolute measure of instrument performance.

| Instrument model | Submitted datasets | Datasets retained after MoltenProt QC | Datasets retained (%) |
| --- | --- | --- | --- |
| Aunty | 5 | 5 | 100 |
| Cary Eclipse | 8 | 2 | 25 |
| Prometheus NT.48 | 73 | 73 | 100 |
| Prometheus NT.Plex | 10 | 10 | 100 |
| Prometheus Panta | 40 | 40 | 100 |
| SUPR-DSF | 20 | 15 | 75 |
| Uncle | 15 | 0 | 0 |
| Total | **171** | **145** | **85** |

**Table S2.** *Effect of fluorescence readout and analysis method on apparent T_m_ for two DSF instruments measured in the same laboratory.* Values are apparent *T*_m_ (°C).

Apparent *T*_m_ values were obtained from measurements obtained with two DSF instruments in the same laboratory under identical sample and acquisition conditions. Fluorescence data were analysed using single‑wavelength intensity at 350 nm, the 350/330 nm ratio, BCM, and SVD, in combination with first derivative or two‑state equilibrium model analysis. BCM and SVD analyses were not available for instrument 15 because this instrument did not record full‑spectrum fluorescence data. Depending on the instrument, fluorescence readout and analysis method, apparent *T*_m_ values differed by up to 7 °C. NA, not available.

| **Instrument** | **Analysis method** | **350 nm** | **350/330 nm ratio** | **BCM** | **SVD** |
| --- | --- | --- | --- | --- | --- |
| **15** | First derivative | 55.2 | 59.2 | NA | NA |
|  | Two‑state equilibrium model | 56.0 | 59.0 | NA | NA |
| **28** | First derivative | 58.5 | 60.0 | 62.5 | 58.5 |
|  | Two‑state equilibrium model | 58.8 | 59.4 | 60.2 | 59.0 |

**Table S3.** *Effect of fluorescence readout and fitting range on apparent ΔH for instrument 28.* Values are mean ± SD (kcal/mol). Instrument 28 was selected for this analysis because its full spectral dataset allowed several fluorescence readouts to be derived from the same measurements: single‑wavelength intensity at 350 nm, the 350/330 nm ratio, BCM and SVD.

| **Fitting range** | **350 nm** | **350/330 nm ratio** | **BCM** | **SVD** |
| --- | --- | --- | --- | --- |
| 40–80 °C | 100.7 ± 4.2 | 101.1 ± 9.8 | 83.4 ± 3.4 | 104.0 ± 3.6 |
| 20–95 °C | 112.0 ± 2.2 | 75.5 ± 7.1 | 63.6 ± 1.3 | 116.5 ± 2.5 |

**Supplementary Protocol**

**Supplementary Protocol S1.**

Protocol distributed to participating laboratories for the MOSBRI interlaboratory differential scanning fluorimetry benchmarking study.

**1. Objective and study design**

The objective of this interlaboratory benchmark was to evaluate the reproducibility of DSF measurements across different instruments and laboratories.

The study was organised in two sequential phases. Participants first received the Phase 1 protocol together with an initial tube of the lyophilised V_H_H reference sample and performed the measurements using their routine acquisition parameters and preferred analysis workflow. After completion and submission of the Phase 1 dataset, participants received the Phase 2 protocol together with a second tube of the same V_H_H reference sample and repeated the measurements using a harmonised acquisition protocol to facilitate direct comparison between laboratories.

**2. Reference sample**

The reference material was a lyophilised V_H_H sample developed through the European MOSBRI project. The V_H_H was pre‑characterised before distribution to participating laboratories. A product information sheet describing the reconstitution procedure was provided with each sample.

**3. Sample preparation**

The lyophilised V_H_H was reconstituted in water according to the product information sheet. After reconstitution, participants were instructed to record a UV‑visible absorbance spectrum between 240 and 340 nm using a standard 1 cm pathlength cuvette, and to determine the protein concentration from the absorbance at 280 nm.

The reconstituted V_H_H was then diluted in PBS buffer to a final concentration of 0.3 mg/mL. The final concentration was verified by UV‑visible spectroscopy before DSF acquisition. Participants were asked to save the absorbance spectra recorded after reconstitution and after dilution.

**4. Phase 1: user-defined DSF protocol**

In Phase 1, each laboratory performed five replicates using its routine acquisition settings and preferred analysis workflow. No restrictions were imposed on the temperature ramp, temperature range, excitation source, fluorescence detection settings or analysis software. This phase was designed to reflect routine laboratory practice as closely as possible.

**5. Phase 2: harmonised DSF protocol**

In Phase 2, all participating laboratories performed five replicate measurements using a common acquisition protocol. The V_H_H sample was adjusted to 0.3 mg/mL in PBS. Measurements were performed using the parameters listed below. Where the exact fluorescence settings were not available on a given instrument, participants used the closest equivalent settings provided by the manufacturer.

| **Parameter** | **Harmonised setting** |
| --- | --- |
| Sample concentration | 0.3 mg/mL in PBS |
| Number of replicates | 5 |
| Temperature range | 20–95 °C |
| Temperature ramp | Linear ramp of 1 °C/min |
| Equilibration time | 5 min before the start of the thermal scan |
| Excitation wavelength | 280 nm, or the closest equivalent instrument setting |
| Emission range | 300–400 nm, or the closest equivalent instrument setting |

**6. Practical precautions**

- Participants were instructed to switch on the instrument at least 15 min before measurement when lamp stabilisation was required.
- Participants were instructed to avoid bubble formation during sample loading.
- For capillary-based instruments, participants were instructed to handle the capillaries only by their extremities and to avoid touching the optical detection area.
- For NanoTemper instruments, participants were instructed to clean the mirror with ethanol before adding capillaries and after completion of the experiment.

**7. Data and metadata requested from participants**

For each phase, participants were asked to provide raw and processed fluorescence data in CSV format, UV‑visible absorbance spectra recorded after reconstitution and after dilution to 0.3 mg/mL, and a brief description of the method or algorithm used to determine apparent melting temperature (*T*_m_) and onset temperature (*T*_onset_) values.

Participants also reported the main acquisition, sample-handling and analysis metadata listed in the reporting template below.

| **Category** | **Information requested** |
| --- | --- |
| Laboratory information | Laboratory name |
| Method information | UV spectrophotometer manufacturer and model |
| Concentration measurement | Concentration after reconstitution in water (mg/mL) |
| Concentration measurement | Concentration after dilution to 0.3 mg/mL |
| Method information | DSF instrument manufacturer and model |
| Method information | Software used for acquisition and analysis |
| Method information | Starting and final temperatures (°C) |
| Method information | Equilibration time before measurement (min) |
| Method information | Excitation and emission settings, or closest equivalent settings |
| Method information | Temperature ramp (°C/min) |
| Experimental metadata | Measurement date |
| Experimental metadata | Time between sample preparation and measurement |
| Summary of results | *T*_m_ values for replicates 1–5 |
| Summary of results | *T*_onset_ values for replicates 1–5 |
| Additional information | Comments on acquisition, sample handling, data processing or deviations from the protocol |
